# Single-cell transcriptomic and neuropathologic analysis reveals dysregulation of the integrated stress response in progressive supranuclear palsy

**DOI:** 10.1101/2023.11.17.567587

**Authors:** Kristen Whitney, Won-Min Song, Abhijeet Sharma, Diana K. Dangoor, Kurt Farrell, Margaret M. Krassner, Hadley W. Ressler, Thomas D. Christie, Ruth H. Walker, Melissa J. Nirenberg, Bin Zhang, Steven J. Frucht, Giulietta M Riboldi, John F. Crary, Ana C. Pereira

**Author notes:** These authors contributed equally to this work (Kristen Whitney, Won-Min Song, Abhijeet Sharma). Co-senior authors (John F. Crary, Ana C. Pereira). **Address correspondence to:** Dr. Ana. C. Pereira, Department of Neurology, Icahn School of Medicine at Mount Sinai, 1468 Madison Avenue, New York, NY, 10029, USA Dr. John. F. Crary, Icahn School of Medicine at Mount Sinai, 1 Gustave L. Levy Place Box 1194, New York, NY 10029.

## Abstract

Progressive supranuclear palsy (PSP) is a sporadic neurodegenerative tauopathy variably affecting brainstem and cortical structures and characterized by tau inclusions in neurons and glia. The precise mechanism whereby these protein aggregates lead to cell death remains unclear. To investigate the contribution of these different cellular abnormalities to PSP pathogenesis, we performed single-nucleus RNA sequencing and analyzed 45,559 high quality nuclei targeting the subthalamic nucleus and adjacent structures from human post-mortem PSP brains with varying degrees of pathology compared to controls. Cell-type specific differential expression and pathway analysis identified both common and discrete changes in numerous pathways previously implicated in PSP and other neurodegenerative disorders. This included EIF2 signaling, an adaptive pathway activated in response to diverse stressors, which was the top activated pathway in vulnerable cell types. Using immunohistochemistry, we found that activated eIF2α was positively correlated with tau pathology burden in vulnerable brain regions. Multiplex immunofluorescence localized activated eIF2α positivity to hyperphosphorylated tau (p-tau) positive neurons and ALDH1L1-positive astrocytes, supporting the increased transcriptomic EIF2 activation observed in these vulnerable cell types. In conclusion, these data provide insights into cell-type-specific pathological changes in PSP and support the hypothesis that failure of adaptive stress pathways play a mechanistic role in the pathogenesis and progression of PSP.

## INTRODUCTION

Abnormal tau accumulation is the common neuropathological feature of a group of neurodegenerative disorders called tauopathies. While Alzheimer’s disease (AD) is the most common tauopathy, progressive supranuclear palsy (PSP) is the most common primary amyloid-independent tauopathy, estimated to affect five in every 100,000 people [1]. On the cellular level, PSP is characterized histopathologically by deposition of abnormal hyperphosphorylated tau (p-tau) in neurons and glia predominantly in the basal ganglia and brainstem [2–6]. Characteristic neuropathologic features are neuronal loss, tau-positive pre-tangles and neurofibrillary tangles (NFTs), neuropil threads, oligodendroglial coiled bodies, and morphologically distinct tufted astrocytes, a hallmark feature for PSP [7, 8]. Tau pathology in oligodendrocytes forming coiled bodies is a feature in PSP and other tauopathies, including corticobasal degeneration (CBD), argyrophillic grain disease (AGD) and globular glial tauopathy (GGT) [9–11]. Recently published PSP neuropathological diagnostic criteria includes neurofibrillary tangles in the globus pallidus, substantia nigra, and subthalamic nucleus, and tufted astrocytes in the peri-Rolandic cortices and putamen [7]. Multiple PSP clinical subtypes and stages are distinguished by neuroanatomic distribution of tau pathology [12]. Across subtypes, the subthalamic nucleus is one of the first, and the most severely affected brain regions in this disease [13]. Mechanisms underlying PSP progression are unclear [12]. Understanding the molecular changes occurring in vulnerable cell types is important to identify candidate mechanistic drivers of PSP pathology, novel therapeutic targets, and both disease-specific pathways as well as common pathogenic mechanisms of tau-mediated neurodegeneration.

PSP is largely a sporadic disease, with rare reported cases of autosomal dominant mutations in the *MAPT* gene causing a PSP-like syndrome [14–16]. Genome wide association studies (GWAS) have identified a common haplotype in the 17q21.31 *MAPT* locus as the major genetic risk factor [17–19]. The relationship between this genetic risk and disease pathogenesis remains unclear but likely stems from differences in *MAPT* expression and splicing. Several PSP-associated risk alleles outside of the *MAPT* locus have been identified, including single nucleotide polymorphisms (SNPs) near *STX6*, *MOBP* and *EIF2AK3* [17–19]. *EIF2AK3* encodes the endoplasmic reticulum (ER) membrane protein, protein kinase R-like ER kinase (PERK), a stress sensor in the unfolded protein response (UPR), which maintains cellular homeostasis under conditions of protein folding disturbances [20]. PERK activation phosphorylates eukaryotic translation initiation factor 2 alpha (eIF2α), which is a key component of the integrated stress response (ISR), a common adaptive pathway for restoring cellular homeostasis by inhibiting mRNA translation and protein synthesis [21]. The UPR has been shown to be involved in multiple neurodegenerative diseases including PSP [22–26]. Previous bulk transcriptomic analyses using cortical and cerebellar autopsy PSP brain tissue identified transcriptional changes that potentially drive distinct cell-specific tau aggregation [27]. This dataset also identified a positive association of NFTs with synaptic genes and tufted astrocytes with microglial gene-enriched immune networks. This study implicates diverse molecular mechanisms underlying cell-type specific vulnerability to tau aggregation and degeneration. However, the extent to which this represents differences in the relative abundance of cells or cell-type specific gene expression changes is challenging to determine with bulk transcriptomic data.

Recent studies in post-mortem brain tissue using single-cell sequencing have begun to elucidate cell-type specific changes in different neurodegenerative diseases including AD, Parkinson’s disease (PD) and Huntington’s disease (HD) [28–34]. Here, we characterized the transcriptional changes and cellular diversity of the subthalamic nucleus and surrounding regions in PSP and clinically normal control brains using droplet based single-nucleus RNA sequencing (snRNA-seq), a massively parallel single-nucleus sequencing method. We focused our analysis on neurons, astrocytes and oligodendrocytes, the cell types most vulnerable to toxic tau accumulation in PSP, although significant gene expression differences were identified in all cell populations, including a novel astrocyte-oligodendrocyte hybrid population. We observed unique and shared expression profiles in vulnerable cell types, identified EIF2 signaling as the top dysregulated pathway, and histologically validated eIF2α activation in neurofibrillary and glial tau inclusions in PSP vulnerable brain regions. Our findings support the hypothesis that adaptive stress pathways play a mechanistic role in PSP pathogenesis and highlight the importance of combining single-nucleus molecular data with rigorous neuropathological target validation.

## METHODS

### Human brain tissue samples, nuclei isolation and single-nucleus RNA sequencing

Postmortem fresh frozen autopsy brain tissue and formalin-fixed paraffin-embedded (FFPE) brain tissue from control and autopsy-confirmed PSP cases were obtained from the Neuropathology Brain Bank & Research CoRE at Mount Sinai (New York, NY). Subjects were selected based on their final neuropathological diagnosis at autopsy and were matched according to age, sex, and post-mortem interval (PMI). All controls were cognitively normal and negative for established neurodegenerative proteinopathies at the time of autopsy. Frozen tissue blocks containing the subthalamic nucleus and adjacent structures were mounted on a cryostat and 30µm scrolls were cut to ensure equal representation of the entire block. Nuclei were isolated from the scrolls immediately. Tissue scrolls were homogenized in a dounce homogenizer with Nuclei EZ lysis solution. The suspension from each sample was filtered through a 40um Flowmi^TM^ strainer (Bel-Art), washed, re-filtered, and resuspended in 0.05% BSA PBS containing RNAsin RNAse inhibitor. Nuclear suspension was counted and processed by the Chromium Controller (10x Genomics) using single Cell 3′ Reagent Kit v2 (Chromium Single Cell 3′ Library & Gel Bead Kit v2, catalog number: 120237; Chromium Single Cell A Chip Kit, 48 runs, catalog number: 120236; 10x Genomics). Single-nucleus RNA-sequencing (snRNA-seq) was performed by Genewiz.

### SnRNA-seq quality control (QC)

SnRNA-seq data was QCed (snQC) by employing *scran* package (v 1.14.6) [35]. Low-quality cells were removed by outlier detection for low library size and proportion of mapped endogenous reads with Median Absolute Deviation (MAD) > 3. To further filter out low quality cells, the metric mitochondrial read rate (T_mito_) was applied as a proportion of mitochondrial transcripts in each cell’s library. Adaptive mitochondrial rate thresholds were applied to remove highly apoptotic cells while minimizing the loss on functional transcripts. The number of expressed genes in cells (as a proxy for functional information) was analyzed across a range of mitochondrial rate thresholds (T_mito_) between 10-70%. Per threshold, the representative number of expressed genes (N_eg_) was obtained as the lower quantile value among the filtered cells. As the lower threshold enriches for cells with better quality, it is expected that the overall number of expressed genes abruptly increases. On the other hand, the higher threshold will enforce inclusion of cells with lower number of expressed genes, thus N_eg_ will gradually reach its plateau towards the N_eg_ of all cells without any mitochondrial rate thresholding. To identify the balance between these two extremities, adaptive thresholds were obtained from the elbow in the T_mito_ -vs-N_eg_ curve. Per sample, doublet cells were evaluated by DoubletFinder() (v 2.0.3) from *Seurat* and doubletCells() from *scran* [36]. Cell clusters enriched with high confidence calls from DoubletFinder (FET FDR < 0.05) and high doublet scores from doubetCells (T-test FDR < 0.05) were removed. Individual cells called with high-confidence doublets by DoubletFinder were also removed, and the dropout reads were estimated using SAVER [37].

### Data merging and batch correction

The QCed data from individual samples were combined and corrected for batch effects using Mutual Nearest Neighbor (MNN) in the *batchelor* R package (v1.2.4) [38]. To mitigate differences in variance between batches, cells were down-scaled to match the least-sequenced batch’s coverage using “multiBatchNorm()” in *batchelor* package [39]. Technical and biological variances in individual gene expressions within each batch were modeled using ‘modelGeneVar()’ from the scran R package (v1.16.0). The technical variations were inferred by fitting variance-mean curve as the technical variation, and the deviation from the fitted variance-mean curve was modeled as biological variations with ‘modelGeneVar()’ from the scran R package (v1.16.0). Variances were combined across the batches with ‘combineVar()’ function from the scran R package (v1.16.0), and extracted variable genes exhibiting any biological variance. The MNN-based batch correction was then executed by “fastMNN()” function from *batchelor* package on the chosen variable genes set.

## 2-Step Unsupervised cell clustering

For the first step of cell clustering, k-Nearest-Neighbor (kNN) graph-based clustering was performed within batch corrected feature space and MNN in *batchelor* package. To obtain the optimal kNN parameter to construct kNN graph, walktrap clustering was performed on each kNN ɛ [log(N_c_),√N_c_] where N_c_ = number of cells. The ‘cluster_walktrap()’ function was implemented in *igraph* package (v 1.2.5) [40]. The clustering solutions across different kNN were then compared to each other by adjusted Rand index to evaluate degree of similarity. kNN=55 showed the highest overall similarity to the other solution (i.e., the centroid), and was chosen as the first-step solution with 44 cell clusters (**Supplemental figure 2b-c**). While this first-step solution provided fine-resolutions to the cell populations, we sought to identify coarse-grained clusters reflecting more robust cell populations. For the second cell clustering step, inter-cluster similarity was evaluated to merge similar cell clusters. Correlation distance d_ij_ = √(2(1-ρ_ij_) was utilized as the distance metric between the clusters, where ρ_ij_ is Pearson correlation between two cells, i and j, within MNN-corrected feature space, as the cell-cell distance. Then, inter-cluster distance between clusters a and b, D_ab_, was calculated by D_ab_ = d[a,b]/[r(a) + r(b)], where d[a,b] = distance between centroid cells in clusters a and b, r(a), r(b) = radius of cluster a or b. r(a) was defined as the distance between the centroid cell of cluster a, and 90% quantile distance to other cells in the same cluster. Overall, D_ab_ represents a separation ratio between two cell clusters, where D_ab_ > 1 if the centroids are farther apart than the radii of the clusters, D_ab_ < 1 otherwise. Based on the separation metric, hierarchical clustering was performed among the clusters with average linkage method (**Supplemental figure 2d**), and the cutoff evaluated by inspecting the overall separation ratio (i.e., average of pairwise D_ab_) against the minimum separation ratio (**Supplemental figure 2e**). The elbow point chosen corresponds to average D_ab_= 0.6478 (**Supplemental figure 2e**). This yielded 13 clusters with distinct sample compositions (**Supplemental figure 2f**).

### De novo cluster marker detection

As cell clusters represent distinct cellular transcriptional states, they can be characterized by distinctively over-expressed genes in each cluster in comparison to other cells, hence *de novo* cluster markers. To this end, we performed pairwise differential expression analysis among the 13 cell clusters by utilizing “*findMarkers()*” function in *scran* (v1.24.0) R package. findMarkers() first performs student t-test for all pairs of cell clusters with normalized, log-transformed expressions, then summarizes the overall p-values and FDR by averaging over the log-transformed p-values, followed by Benjamini-Hochberg correction. FDR < 0.05 and fold-change > 1.2 was applied to identify differentially expressed genes in each cluster.

### Cluster marker analysis and cell type classification

Classes for the 13 cell clusters were assigned to six major cell types (neurons, astrocytes, oligodendrocytes, oligodendrocyte precursor cells (OPCs), microglia and endothelial cells) using the following marker genes: *SLC17A6* and *NRGN* for excitatory neurons; *GAD1 and GAD2* for inhibitory neurons; *AQP4, GFAP,* and *SLC1A3* for astrocytes; *PLP1, MBP* and *MOG* for oligodendrocytes; *VCAN* and *PDGFRA* for OPCs; *CX3CR1, TYROBP,* and *CD74* for microglia; and *FLT1* and *CLDN5* for endothelial cells [29, 32]. For the two clusters expressing markers for either none or more than one major cell type, top marker genes were manually examined and compared to cell-type specific gene lists generated from the literature [30, 41–44]. The cluster characterized by high proportions of both astrocyte and oligodendrocyte lineage genes were defined as hybrid cells. This resulted in two additional cell classes – hybrid cells and choroid plexus (ChP) epithelial cells. Finally, we performed GO term enrichment analysis on the top cluster marker genes and examined the top enriched functions and pathways.

### Cell cluster-specific differential expression between PSP and healthy control cells

For each cell cluster, we performed differential expression analysis, comparing cells from PSP samples to the healthy control samples. First, we normalized the gene expression data using SCTransform, then adjusted the normalized data using sequencing depth as covariates with PrepSCTFindMarkers() function in Seurat (v4.1.1) R package [45]. Then, we performed MAST analysis to compare PSP cells against the healthy control cells within each cell cluster [46]. The significant differentially expressed genes (DEGs) were identified with FDR < 0.05. Further, we required the up-regulated DEGs to be expressed in a higher frequency within the PSP cells than the control cells, and down-regulated DEGs to be expressed in a lower frequency within the PSP cells than the control cells. Finally, differentially regulated pathway and disease and function activation state analysis was performed with Ingenuity Pathway Analysis (IPA, Qiagen). The average log (fold change) and adjusted p-value of each significant (FDR < 0.05) differentially expressed gene was supplied to IPA.

### Cluster specific risk allele expression

PSP GWAS risk genes were identified for downstream analysis [17–19]. After obtaining the normalized, log-transformed expressions, overall p-values and FDR between PSP cases and controls for each cluster, we overlapped these genes with the ten risk genes and visualized the absolute log fold change plotted for each cluster cell type.

### Immunohistochemistry

Formalin-fixed paraffin-embedded (FFPE) tissue sections (5 μm) from blocks containing the basal ganglia (i.e., substantia nigra, putamen, globus pallidus, caudate, subthalamic nucleus), thalamus, Peri-Rolandic cortices, cerebellum containing the dentate nucleus and visual cortex were mounted onto positively charged slides, and baked overnight at 70°C. Sections were stained with Luxol fast blue counterstained with hematoxylin & eosin (LHE) on a Leica X Autostainer (Leica Biosystems, Wetzlar, Germany). Immunohistochemistry (IHC) using an antibody targeting phosphorylated tau (AT8, pSer202/pThr205, 1:1000, Thermo Scientific MN1020) was performed on the Ventana Benchmark XT and Ventana Discovery Ultra automatic staining platforms (Ventana Medical Systems, Oro Valley, AZ) according to manufacturer’s instructions with reagents and antibodies acquired from the same lot and positive and negative batch controls. Sections were boiled in CC1 (citric acid buffer, Roche Diagnostics, Basel Switzerland) for 1 hour (pH 6) for antigen retrieval, followed by primary antibody incubation for 36 minutes and visualized using the Ultraview detection kit (Roche Diagnostics) followed by 3,3’-diaminobenzidine (DAB). IHC using antibodies targeting phosphorylated peIF2α (pSer51, 1:1000, Millipore Sigma SAB4504388), and phosphorylated tau (AT8, pSer202/pThr205, 1:1000, Thermo Scientific MN1020) Biosystems, Wetzlar, Germany) was performed on the Leica Bond Rx automatic staining platform according to manufacturer’s instructions with reagents and antibodies acquired from the same lot and positive and negative batch controls. Heat-induced epitope retrieval (HIER) was performed with EDTA-based pH 6 epitope retrieval solution (Leica Biosystems, Wetzlar, Germany) followed by primary antibody incubation for 20 minutes and DAB for visualization. For slides that were double labeled, both DAB and alkaline phosphatase were used for visualization. All slides were counterstained with hematoxylin and cover slipped. Slides were imaged using an Aperio Versa 8 (Leica Biosystems, Wetzlar, Germany) and NanoZoomer S210 (Hamamatsu, Shizuoka, Japan) digital slide scanner.

### Semiquantitative and digital quantitative histopathologic assessments

All cases were analyzed using Aperio ImageScope software (versions 9 and 12.3, Leica Biosystems). Neuroanatomical regions were manually segmented on LHE-stained whole slide images and transferred to p-tau (AT8) and peIF2α immunostained sections. For each brain region except the subthalamic nucleus, a 4 mm^2^ region was annotated within each neuroanatomical region. Due to the elongated shape and disease-related atrophy of the subthalamic nucleus, the entire region was annotated and normalized to mm^2^. Hyperphosphorylated tau (AT8) burden was semi-quantitatively scored from 0-3 (0 = none, 1 = rare, 2 = moderate, 3 = severe) for neuropil threads, neuronal, and glial tau burden according to the Braak criteria [47]. For digital quantitative assessment of tau burden, the Aperio positive pixel count algorithm (version 9) was run in the same 4 mm^2^ region using DAB intensity parameters set on the batch control. Total positive pixel values were automatically generated and normalized to total number of pixels detected. For peIF2α quantification, in each brain region the number of peIF2α-positive cells was manually counted in the 4 mm^2^ annotation or the entire subthalamic nucleus and normalized to the number of positive cells per mm^2^. Phospo-eIF2α counts were scaled to semiquantitative scores from 0 (-) to 3 (+++), 0 = no positive cells, 1 = less than 1 positive cell/mm^2^, 2 = less than 2 positive cell/mm^2^, 3 = more than 2 positive cell/mm^2^.

### Immunofluorescence microscopy

Multiplex immunofluorescence using antibodies targeting peIF2α (pSer51, Millipore Sigma SAB4504388), phosphorylated tau (AT8, pSer202/pThr205, Thermo Scientific MN1020), ALDH1L1 (Abcam, ab190298), and DAPI was performed on the Leica Bond Rx according to manufacturer’s instructions. Secondary antibodies conjugated to Alexa Fluor 488, 594, and 647 fluorophores were used. Slides were imaged on a Zeiss AxioImager Z2M.

### Statistics

Phospo-eIF2α counts across neuroanatomical regions were compared between PSP and control groups using Fishers exact test (FET). Correlations between peIF2α, semiquantitative tau burden and positive pixel tau burden were analyzed using Pearson’s test. All analysis was performed using GraphPad Prism (version 9.4.0, La Jolla, CA) with significant p value set at < 0.05.

## Results

### Single-nucleus analysis identifies major and novel cell populations

To investigate the cellular diversity and cell-type specific disease-related gene expression changes occurring in PSP, we isolated nuclei from the subthalamic nucleus and surrounding regions from three PSP cases and three age-matched controls (**Table 1**). PSP disease stage was assessed by semiquantitative evaluation of immunohistochemically stained formalin fixed paraffin embedded (FFPE) tissue from the contralateral subthalamic nucleus and adjacent regions using antisera recognizing hyperphosphorylated tau (p-tau) at Ser202/Thr205 (AT8). Staining confirmed neuropathological diagnosis with varying tau burdens in the PSP cases, spanning the progression of PSP-type tau pathology (**Supplemental Figure 1**). Semi-quantitative assessment of disease severity measured neurofibrillary tangles (NFTs), neuropil threads (NT), and glial fibrillary tangles (GFT) including tufted astrocytes and coiled bodies in the PSP cases (**Table 1**). PSP3 had the overall highest tau burden, with severe NFT, NT, and GFT burden representing the late stage of disease progression. Both PSP2 and PSP3 had a higher glial tau burden than PSP1, which had an overall lower tau burden and was likely in the early stage of disease.

**Table 1.** SnRNA-seq Patient data.

| Case # | Sex | Age of Death (yr) | PMI (hr) | Brain weight (g) | Clinical Diagnosis | Neuropath Category | STN |  |  | Thalamus |  |  |
| --- | --- | --- | --- | --- | --- | --- | --- | --- | --- | --- | --- | --- |
|  |  |  |  |  |  |  | NT | NFT | GFT | NT | NFT | GFT |
| PSP1 | M | 64 | 20.8 | 1400 | None | PSP | + | + | + | + | + | + |
| PSP2 | M | 69 | 15.3 | 1364 | PSP | PSP | + | + | ++ | + | + | ++ |
| PSP3 | M | 71 | 11.7 | 1374 | PSP | PSP | +++ | +++ | +++ | + | ++ | ++ |
| Control 1 | M | 73 | 25.4 | 1380 | None | Control |  |  |  |  |  |  |
| Control 2 | M | 71 | 43.8 | 1380 | None | Control |  |  |  |  |  |  |
| Control 3 | M | 67 | 15.4 | 1300 | None | Control |  |  |  |  |  |  |

PMI = postmortem interval (death to freezing)
Semiquantitative scores (0 = none, + = mild, ++ = moderate, +++ = severe)
NT = neuropil threads; NFT = neurofibrillary tangles, GFT = glial fibrillary tangles

Droplet–based single-nucleus RNA-sequencing (snRNA-seq) was performed on the six samples using the 10X Genomics platform (**Figure 1a**). Quality control (QC) measures showed that two PSP samples had preferentially higher mitochondrial rates than the controls and retained less than 50% of all cells with the conventional < 10% threshold. As PSP is a neurodegenerative disease with a greater proportion of apoptotic cells than healthy control samples, we applied an adaptive mitochondrial rate threshold that removes highly apoptotic cells while minimizing the loss on functional transcripts to balance the number of detected genes (**Supplemental Figure 2a**). Further filtering for low depths and imputing for dropout reads resulted in 45,559 high quality nuclei profiles with a median 1,041 genes detected per nucleus. Of these, 18,727 were PSP nuclei and 26,832 were control nuclei (**Table 2**). The overall numbers of QCed cells show a lower proportion of cells from PSP samples remained than controls.

**Figure 1.**
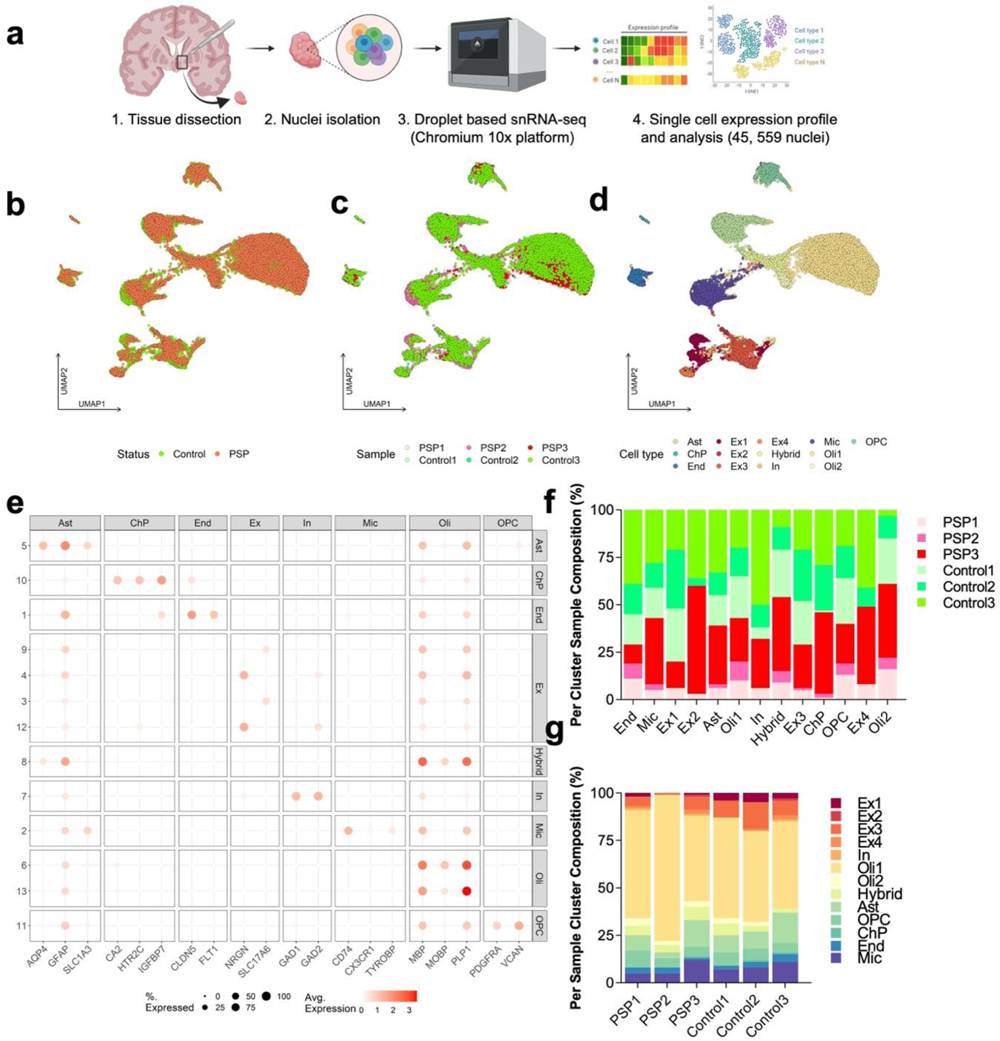
Single-nuclei sequencing of the subthalamic nucleus and adjacent structures in control and PSP post-mortem brain identifies cell-type specific marker genes. **(a)** Schematic of experimental design and snRNA-seq workflow. **(b-d)** UMAP projection of cell populations colored by **(b)** disease diagnosis, **(c)** individual of origin, and **(d)** major cell type. The major cell types identified are oligodendrocytes (Oli1, Oli2), microglia (Mic), astrocytes (Ast), endothelial cells (End), neurons marked as excitatory (Ex1, Ex2, Ex3, Ex4) or inhibitory (In), and oligodendrocyte precursor cells (OPC). Two additional cell types are marked as hybrid and choroid-plexus (ChP). **(e)** Marker gene expression in subclusters. Dot size depicts proportion of cells in each cluster expressing the marker gene, and heat map depicts enrichment of marker expressing cells in the cluster. **(f)** Relative proportion of each cell cluster belonging to individual samples. **(g)** Relative proportion of per sample cluster composition.

**Table 2.** SnRNA-seq QC Summary.

| Sample ID | Before QC |  | After QC |  | Summary |  |  |
| --- | --- | --- | --- | --- | --- | --- | --- |
|  | Genes | Cells | Genes | Cells | Mito. Rate threshold | Median # Genes | QC pass rates (%) |
| PSP 1 | 33538 | 6004 | 18543 | 3829 | 38 | 1020 | 63.77 |
| PSP 2 | 33538 | 4221 | 16762 | 3070 | 34 | 823 | 72.73 |
| PSP 3 | 33538 | 13699 | 21696 | 11828 | 27 | 1338 | 86.34 |
| Control 1 | 33538 | 10907 | 20230 | 9310 | 33 | 979 | 85.36 |
| Control 2 | 33538 | 8516 | 20181 | 7256 | 34 | 1056 | 85.24 |
| Control 3 | 33538 | 11931 | 20526 | 10266 | 25 | 898 | 86.04 |

Nuclei expression profiles were clustered using a two-step unsupervised method based on k-nearest neighbor approach and mapped to individual cell types. The first-step solution provides fine-resolutions to the cell population architecture, while the second step evaluates inter-cluster similarity to identify coarse-grained clusters reflecting more robust cell populations (**Supplemental Figure 2b-e**). The second step of unsupervised clustering yielded 13 cell clusters (**Supplemental Figure 2f**). Separating the cells by disease status and individual samples showed that PSP cells both overlap with and segregate from controls across the 13 clusters (**Figure 1b-c**).

To identify cell type and understand the molecular features and functions that define the clusters, we assigned distinct marker genes for each cluster, compared over-expressed genes within clusters (FDR < 0.05 and fold-change > 1.2), and examined the top enriched functions and pathways by gene ontology (GO) analysis (**Supplemental Table 1**). All major cell types (excitatory neurons, inhibitory neurons, astrocytes, oligodendrocytes, microglia, oligodendrocyte precursor cells (OPC), and endothelial cells) were identified within the 13 clusters based on previously established cell-type-specific marker genes [30, 32, 41–43] (**Figure 1d**). GO enrichment analysis confirmed the enriched functions and pathways of the top cluster marker genes were consistent with the biological functions of the assigned major cell type (**Supplemental Figure 2g**). For example, astrocytes expressed canonical marker genes as well as other genes related to astrocyte-specific functions, including the gap junction-related gene *GJA1*, and *SLC1A2*, a gene regulating homeostatic astrocyte function.

In addition to all major cell types, we also identified two novel clusters (cluster 8 and cluster 10) (**Supplemental Figure 2f)**. Cluster 8 highly expressed both oligodendrocyte (*PLP1*, *MBP*, *MOBP*) and astrocyte (*GFAP*, *AQP4*, *S100B*) markers genes simultaneously, and were designated as hybrid cells (**Figure 1e**, **Table 3**) [30, 41–43]. GO terms for the hybrid cluster marker genes were enriched for peptide chain elongation and myelin sheath (**Supplemental Figure 2g**). Cluster 10 was marked by several choroid plexus epithelial cell genes including *CA2*, *HTR2C*, and *IGFBP7*, and GO terms for cluster marker genes were enriched for extracellular space, therefore this cluster was designated as choroid plexus (ChP) epithelial cells [44] (**Figure 1e**, **Supplemental Figure 2g**). As expected, the ChP cluster was also marked by several molecules in cerebrospinal fluid (CSF) secretome such as *SERPINF1*, *AQP1*, *ENPP2*, *CLU,* and *NPC2* (**Table 3**). Overall, we obtained 1125 endothelial cells, 4206 microglia, 6306 neurons, 5117 astrocytes, 23,901 oligodendrocytes, 2,610 OPCs, 2069 hybrid cells, and 235 choroid plexus cells.

**Table 3.**
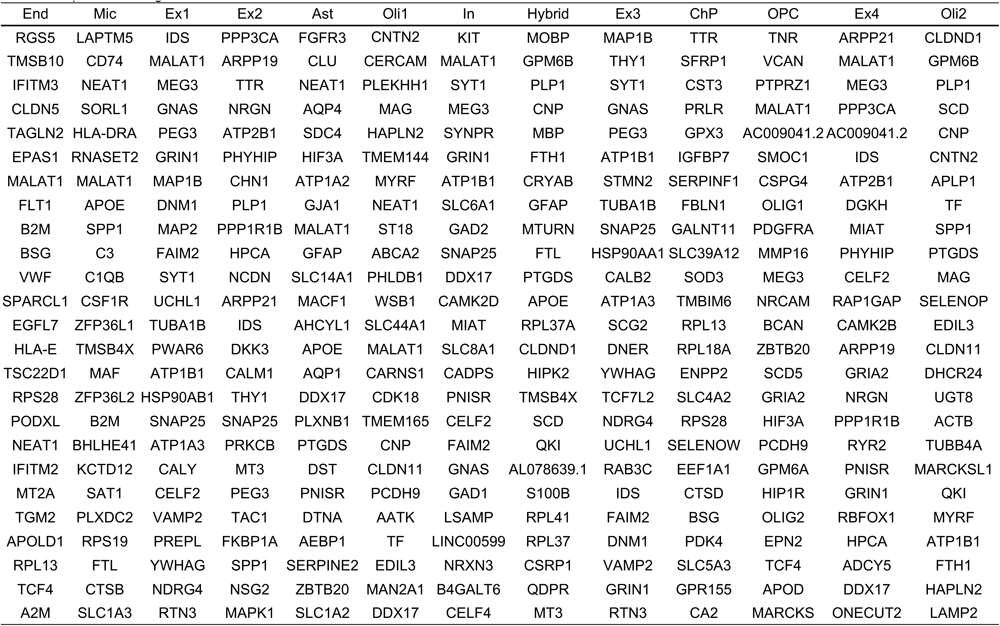
Top 25 marker genes for cell clusters.

To assess whether the proportions of major cell types change with disease state, we compared the relative proportion of each cluster belonging to individual PSP and control samples and the per sample cluster composition (**Figure 1f-g**). Relative cluster sample composition comparison revealed some cell types are under-represented in PSP samples. In non-neuronal cell populations, the proportion of microglia (PSP 43%, control 57%), astrocytes (PSP 39%, control 61%), oligodendrocytes (Oli1 PSP 43%, control 57%; Oli2 PSP 61%, control 39%), OPCs (PSP 40%, control 60%), hybrid (PSP 54%, control 46%) and ChP (PSP 46%, control 54%) were comparable in PSP cases and controls but there was lower relative abundance of endothelial cells (PSP 29%, control 71%) in PSP (**Figure 1f**). In the neuronal populations, abundance of two excitatory neuronal populations, Ex2 and Ex4, were comparable in PSP and controls (Ex2, PSP 61%, control 39%; Ex4. PSP 49%, control 51%), although these proportions were mostly driven by the two samples with the largest threshold adaptive mitochondrial rate cell counts (**Figure 1f**, **Table 2**). These Ex2 and Ex4 neuronal clusters with no apparent neurodegeneration in PSP overexpress *PPP3CA* (PP2B), one of the major phosphatases that directly regulates tau phosphorylation, and *ARPP21* and *ARPP19*, genes that mediate protein phosphorylation through cAMP [48–50] (**Table 3**). In contrast, there was low relative abundance in PSP of two excitatory neuronal populations, Ex1 and Ex3, and the inhibitory neuronal population, In (Ex1, PSP 20%, control 80%; Ex3, PSP 29%, control 71%; In, PSP 33%, control 67%). The excitatory neuronal clusters (Ex1 and Ex3) with potential neurodegeneration in PSP overexpress *CELF2* and *PEG3*, genes which have been shown to induce apoptosis and neuronal death [51–53]. Comparison of the per sample cluster composition shows that across all samples, the majority of cells were glial cells, and particularly oligodendrocytes were the most abundant cell type (**Figure 1g**). This is consistent with the neuroanatomical region targeted, which included a high proportion of white matter (**Supplemental Figure 1a**).

### Differential gene expression analysis identifies dysregulated cellular functions in PSP neurons

Differentially expressed genes (DEG) were examined by cell cluster, then the major pathways, diseases, and functions represented by DEGs were identified and compared between PSP and controls using integrated pathway analysis (IPA). We first focused on differential gene expression between PSP cases and controls in neurons. Across neuronal clusters, 343 genes were downregulated in PSP neurons while 340 genes were upregulated, with the largest differential gene expression in excitatory neuron clusters Ex1 and Ex3 (**Figure 2a**).

**Figure 2.**
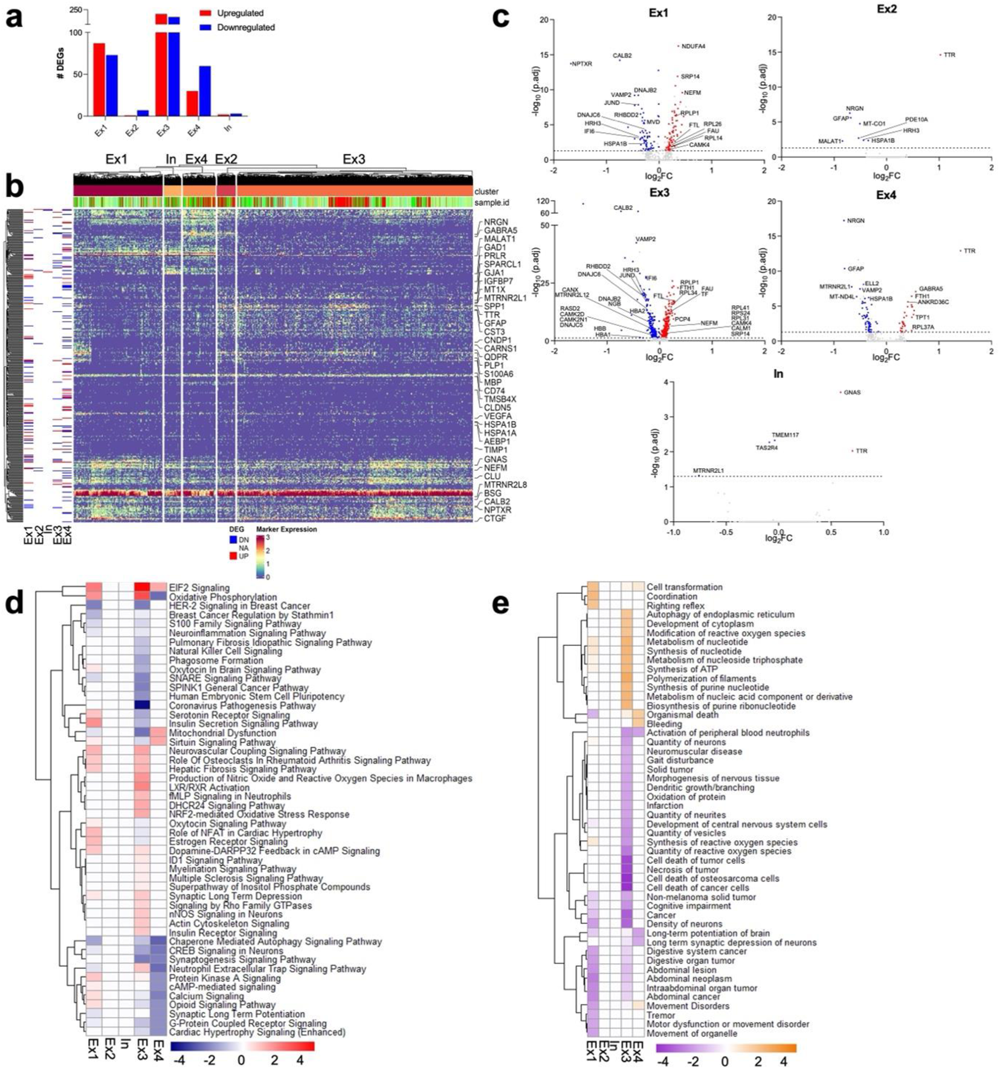
Differentially expressed genes (DEGs) and regulated pathways in neuronal clusters. **(a)** Total numbers of significant upregulated and downregulated genes in excitatory (Ex1, Ex2, Ex3, Ex4) and inhibitory (In) neuron populations between PSP and controls. **(b)** Hierarchical clustering and gene expression heat map of log fold change of top DEGs (log fold change >1.2 and FDR < 0.05) between PSP and control cells in neuronal clusters. **(c)** Volcano plots showing select significant DEGs in individual neuronal clusters. **(d)** Top 50 differentially regulated pathways in PSP neuron clusters derived from ingenuity pathway analysis (IPA). **(e)** Top 50 differentially predicted disease and function activation states in PSP neuron clusters derived from IPA. For (d) and (e), p-values were generated from Fisher’s exact test and heat map is colored by z-score. Positive scores predict an overall increase in pathway activity or disease and function activation states and negative scores predict an overall decrease.

The neuronal clusters that were less abundant in PSP (Ex1, Ex3, and In) were enriched for genes related to apoptosis, neuronal injury, and several genes and pathways critical to neuronal signaling and function. These clusters downregulated multiple genes related to apoptosis regulation, including *JUND*, *IFI6, MTRNR2L12,* and *TMEM117*, and overexpressed *NEFM*, which encodes for a neurofilament protein that is a biomarker of neuronal injury and neurodegeneration [54] (**Figure 2b-c**). Genes related to neurotransmitter release, neuronal arborization and morphology, such as *HRH3, VAMP2*, and *GPRIN3*, were also downregulated in these clusters [55, 56] (**Figure 2b-c**). Several genes related to calcium signaling were significantly differentially expressed such as calmodulin-dependent protein kinase II (*CAMK2A)*, calmodulin (*CALM1)*, and *CALB2,* as well as purkinje cell protein 4 (*PCP4)*, which regulates calmodulin-mediated signaling and neuronal survival [57] (**Figure 2b-c**). Notably, calcium signaling, protein kinase A signaling, and dopamine-DARPP32 feedback in cAMP signaling pathway, a component of an important mechanism regulating glutamatergic and dopaminergic signaling, were also dysregulated in Ex1 and Ex3 [58] (**Figure 2d**). Ex1 and Ex3 were predicted to have lower density of neurons and reduction of several critical functions including decreased dendritic growth or branching, reduced synaptic long-term potentiation, synaptogenesis signaling, and synaptic long-term depression (**Figure 2d-e**). Ex1 was also enhanced for predicted motor dysfunction or movement disorders disease state (**Figure 2e**).

Several genes related to protein translation, topologically incorrect protein degradation, the unfolded protein response, and endoplasmic reticulum (ER) stress including ribosomal, heat shock and chaperone genes were differentially expressed in multiple excitatory neuronal clusters such as *FAU*, *CANX, DNAJB2*, *MVD*, *RHBDD2, HSPA1B,* and *SRP14* (**Figure 2b-c**). Increased EIF2 signaling, a target of the ISR and UPR homeostatic stress response, was the top enriched pathway across neuronal clusters. PSP excitatory neurons also showed reduced chaperone-mediated autophagy signaling (**Figure 2d**).

Additionally, Ex1 and Ex3 neurons overexpressed genes involved in iron homeostasis and heavy metal accumulation, including ferritin light chain (*FTL), FTH1*, and transferrin (*Tf*) which encode for proteins related to ferritin iron storage, and *MT3* which binds heavy metals [59, 60] (**Figure 2b**). Genes related to oxygen homeostasis including alpha and beta hemoglobin (*HBA1*, *HBA2*, *HBB*) and neuroglobin (*NGB*) were significantly downregulated in Ex3 neurons, while NRF2-mediated oxidative stress response was highly enriched in this neuronal cluster (**Figure 2b-d**). Overall, these data provide a summary of the gene expression changes and related functional pathways in PSP neurons, highlighting apoptosis, EIF2 signaling and ER stress, and iron and oxygen homeostasis as dysregulated cellular functions that may underlie disease-related changes seen in PSP neurons.

### Differential gene expression analysis identifies dysregulated cell functions in PSP glial populations

We next investigated the cell-type specific gene expression changes in the glial cells uniquely affected by tau accumulation in PSP, including astrocyte and oligodendrocytes, as well as the novel astrocyte-oligodendrocyte hybrid cell cluster. Our analysis revealed the highest number of DEGs was in PSP oligodendrocytes (Oli1 and Oli2), 1382 genes were upregulated while 656 genes were downregulated. In PSP astrocytes, 1123 genes were upregulated while 217 genes were downregulated, and in the hybrid cluster 48 genes were upregulated while 128 genes were downregulated (**Figure 3a**).

**Figure 3.**
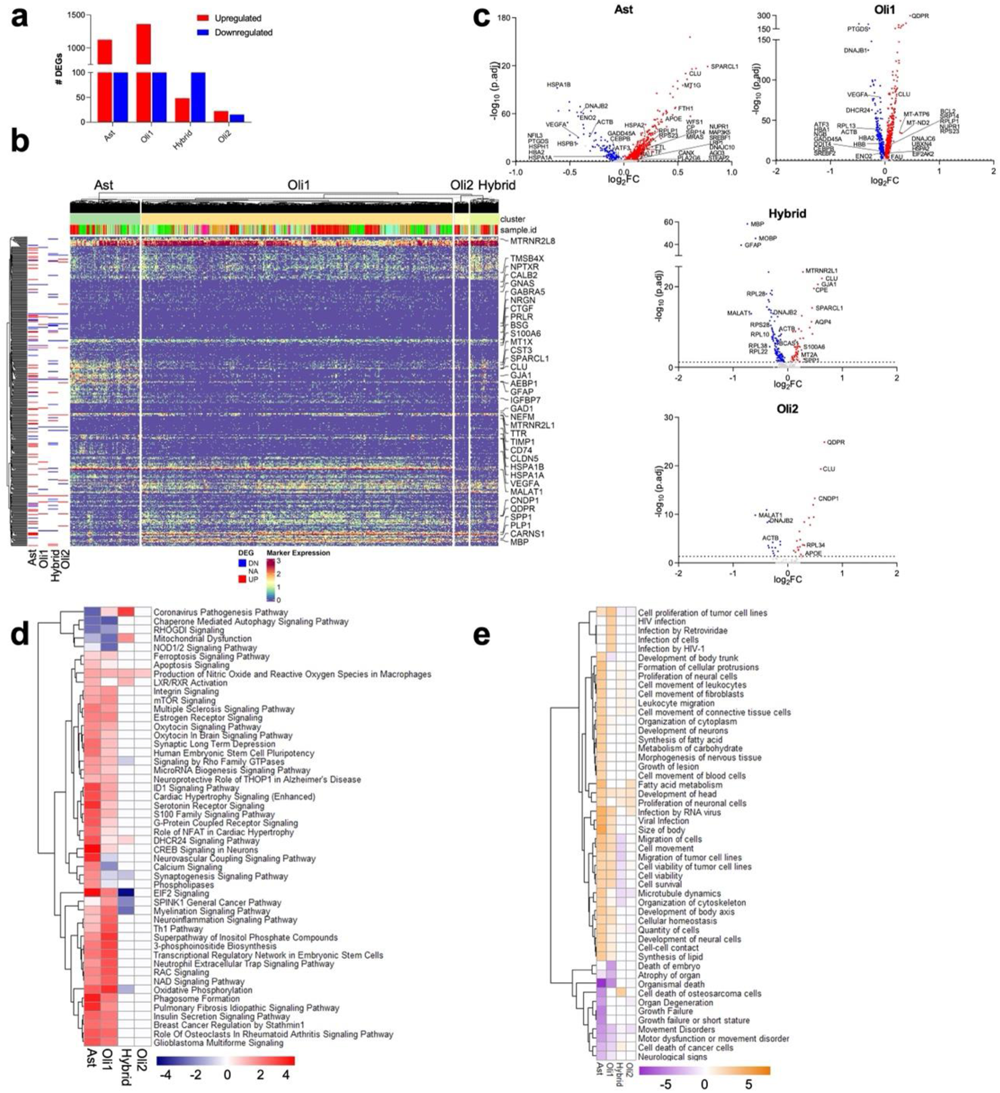
Differentially expressed genes (DEGs) and regulated pathways in astrocyte, oligodendrocyte and hybrid cell populations. **(a)** Total numbers of significant upregulated and downregulated genes in astrocytes (Ast), oligodendrocytes (Oli1 and Oli2), and the hybrid cluster between PSP and controls. **(b)** Hierarchical clustering and gene expression heat map of top DEGs log fold change (log fold change >1.2 and FDR < 0.05) between PSP and control glial clusters. **(c)** Volcano plots showing select significant DEGs in individual glial clusters. **(d)** Top 50 differentially regulated pathways in PSP glia clusters derived from ingenuity pathway analysis (IPA). **(e)** Top 50 differentially predicted disease and function activation states in PSP glial clusters derived from IPA. For (d) and (e), p-values were generated from Fisher’s exact test and heat map is colored by z-score. Positive scores predict an overall increase in pathway activity or disease and function activation states and negative scores predict an overall decrease.

Consistent with the differential gene expression changes and EIF2 activation seen in PSP neurons, several genes related to protein translation, topologically incorrect protein degradation, the unfolded protein response, and ER stress were significantly dysregulated in PSP astrocytes and oligodendrocytes, such as *DNAJB1, DNAJB2, HSPA1A*, *CEBPB*, *HSPB1*, *SREBF1*, *ACTB*, *VEGFA* and *SRP14* as well as several ribosomal genes (**Figure 3b-c**). PSP astrocytes and oligodendrocytes also differentially expressed several UPR genes including PERK-regulated genes *ATF3*, *NUPR1*, *CEBPB*, and *GADD45A,* and were enriched for increased EIF2 signaling [61, 62] (**Figure 3b-d**). In addition to dysregulated stress response, PSP astrocytes and oligodendrocytes also had decreased expression of *ENO2* and *PTGDS,* which code for proteins with potential neuroprotective and anti-apoptotic functions [63, 64] (**Figure 3b-c**). Both PSP astrocytes and oligodendrocytes were also associated with increased mTOR and apoptosis signaling pathways, reduced chaperone-mediated autophagy signaling, and movement disorders and motor dysfunction (**Figure 3d-e**).

Many genes that have been previously linked to other neurodegenerative diseases were also differentially expressed in PSP astrocytes and oligodendrocytes (**Figure 3b-c**). Both PSP astrocytes and oligodendrocytes overexpressed AD-associated genes apolipoprotein E (*ApoE*) and clusterin (*CLU)*, which codes for a chaperone protein involved in several biological events such as cell death [65, 66]. Additional AD-associated genes were differentially expressed in PSP oligodendrocytes, including increased expression of quinoid dihydropteridine reductase (*QDPR)* and decreased expression of *DHCR24*, a gene with cholesterol-synthesizing activity [28, 67]. PSP oligodendrocytes overexpressed *MT-ATP6* and *MT-ND2*, genes which have previously been implicated in Parkinson’s disease and multiple sclerosis [68, 69]. Both PSP astrocytes and oligodendrocytes were also associated with increased activation of pathways related to inflammation and regulation of lipid metabolism, including LXR/RXR signaling and neuroinflammation signaling pathway (**Figure 3d-e**).

In parallel with PSP neurons, iron and oxygen homeostasis was dysregulated in PSP glia. PSP astrocytes overexpressed many genes involved in iron homeostasis, including *CP*, *FTL*, *PLA2G6*, *LRP1*, *Tf*, *STEAP2*, and *FTH*, as well as *MT1G*, which is involved in cellular response to intracellular metal toxicity [59]. Both PSP astrocytes and oligodendrocytes were also associated with increased ferroptosis signaling pathway, and downregulated genes involved in oxygen homeostasis including alpha and beta hemoglobin (*HBA1*, *HBA2*, *HBB*) and neuroglobin (*NGB*) (**Figure 3b-d**).

The PSP astrocyte-oligodendrocyte hybrid cells overexpressed several genes which code for secreted proteins with known neuroprotective functions, including osteopontin (*SPP1*), carboxypeptidase E (*CPE*), and metalothionein2A (*MT2A*) (**Figure 3b-c**). PSP hybrid also overexpressed *S100A6*, an intracellular protein with a regulatory role in several cellular functions including apoptosis, cytoskeleton dynamics, and response to stress [70]. In addition to increased expression of neuroprotective genes, the hybrid cluster was associated with reduced EIF2 signaling (**Figure 3d**). These data provide a summary of the discrete and common gene expression changes and related functional pathways in PSP astrocytes, oligodendrocytes, and the novel astrocyte-oligodendrocyte hybrid cells. We highlight a potential neuroprotective function in the hybrid cells, and EIF2 signaling and ER stress, iron accumulation and oxygen homeostasis, and other pathways previously linked to neurodegeneration, as potential mechanisms which may underlie cell-type specific susceptibility to disease pathology in PSP glia.

### Examination of PSP GWAS candidate risk genes expression across cell types

Large genome wide association studies (GWAS) have identified susceptibility-associated loci and candidate risk alleles in PSP [17–19, 71]. However, which cell types mediate genetic risk and how candidate risk gene expression influences PSP susceptibility is unknown. To investigate this, we examined ten previously identified genes implicated in PSP risk in our snRNA-seq data set by mapping gene expression changes stratified by cell type (**Figure 4**). Strikingly, *MAPT,* the most strongly associated genetic risk factor for PSP, was downregulated in most PSP neuronal clusters except inhibitory neurons (**Figure 4a**). In contrast, we found moderate upregulation in oligodendrocytes (Oli2) with no differential regulation in astrocytes (**Figure 4b**). *EIF2AK3*, which encodes for the UPR stress sensor PERK, was highly expressed in one excitatory neuronal cluster (Ex2) with slight upregulation across all glia clusters. *STX6* was highly expressed in all PSP excitatory neuron clusters but not inhibitory neurons or in any glial cells (**Figure 4a-b**). *STX6* encodes the SNARE protein syntaxin 6 and plays a role in vesicle-membrane fusion in endosomal structures [72]. All PSP neuron clusters, astrocytes and oligodendrocytes (Oli1) highly expressed *SLC01A2* (**Figure 4a-b**). *SLC01A2* is a transporter present at the blood-brain barrier, where it regulates solute trafficking [73]. *MOBP*, a gene previously identified as specific to oligodendrocytes, was downregulated in PSP oligodendrocytes, astrocytes, and hybrid cells as well as in most neuronal clusters (Ex1, Ex2, Ex3, In) (**Figure 4a-b**). Together, these data suggest that PSP candidate risk gene expression may differentially influence selective vulnerability in different cell types and provides further context to understand the complexity of disease gene variants on cell-type specific genetic susceptibility.

**Figure 4.**
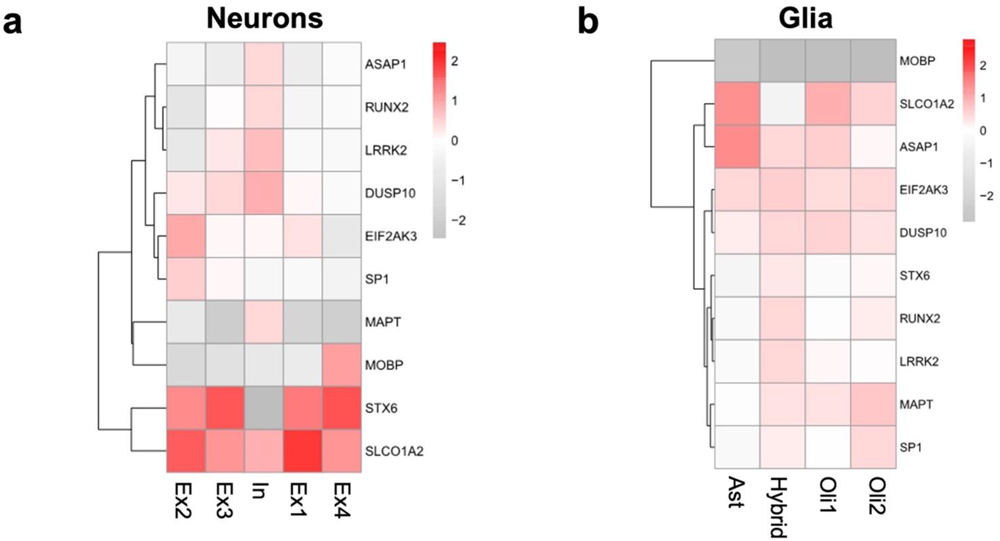
Expression patterns of PSP candidate risk genes across cell types. Heat maps showing the normalized log fold change of differential expression for candidate genes in PSP and control **(a)** neuron clusters, and **(b)** glia clusters (astrocytes, oligodendrocytes, hybrid cells).

### Activated EIF2 signaling in PSP is associated with tau pathology

SnRNA-seq analysis revealed increased EIF2 signaling activation as the top dysregulated pathway in multiple cell populations vulnerable to tau accumulation in PSP, including neurons, astrocytes, and oligodendrocytes. Given that EIF2 is a component of the ISR, an elaborate signaling pathway activated in a range of both normal and pathological contexts, and EIF2 signaling is a target of the ER-stress UPR pathway, we sought to ascertain whether this finding was specifically associated with tau pathology. To investigate this, we performed immunohistochemistry on eight brain regions either selectively vulnerable in PSP (subthalamic nucleus and adjacent thalamus, peri-Rolandic cortex, putamen, globus pallidus, caudate, dentate nucleus of cerebellum) or protected (occipital cortex) on ten PSP cases and six matched-controls using an antibody for phosphorylated eIF2α (peIF2α), the activated form of this protein [7] (**Table 4**). Phospho-eIF2α staining highlighted numerous cytoplasmic granules in positive cells (**Figure 5a**). To quantify peIF2α signal, we counted the number of PSP cases and controls with positive cells in each region (**Figure 5b**). No peIF2α positive (peIF2α^+^) cells were detected in any brain region in the controls. Phospho-eIF2α^+^ cells were detected in the subthalamic nucleus, thalamus, and putamen in nine of the ten PSP cases, and all vulnerable brain regions had significantly more peIF2α^+^ PSP cases than controls (**Figure 5b**). To further investigate peIF2α burden in PSP, we counted the number of peIF2α^+^ cells in the sampled vulnerable and protected PSP brain regions and scored semi-quantitatively from absent (-) to >2 positive cells per mm^2^ (+++) (**Figure 5c**). The putamen, thalamus, subthalamic nucleus, and caudate had the highest number of peIF2α^+^ cells while no peIF2α^+^ cells were detected in the occipital cortex. These data show regional peIF2α burden is consistent with patterns of tau burden across vulnerable brain regions in PSP.

**Figure 5.**
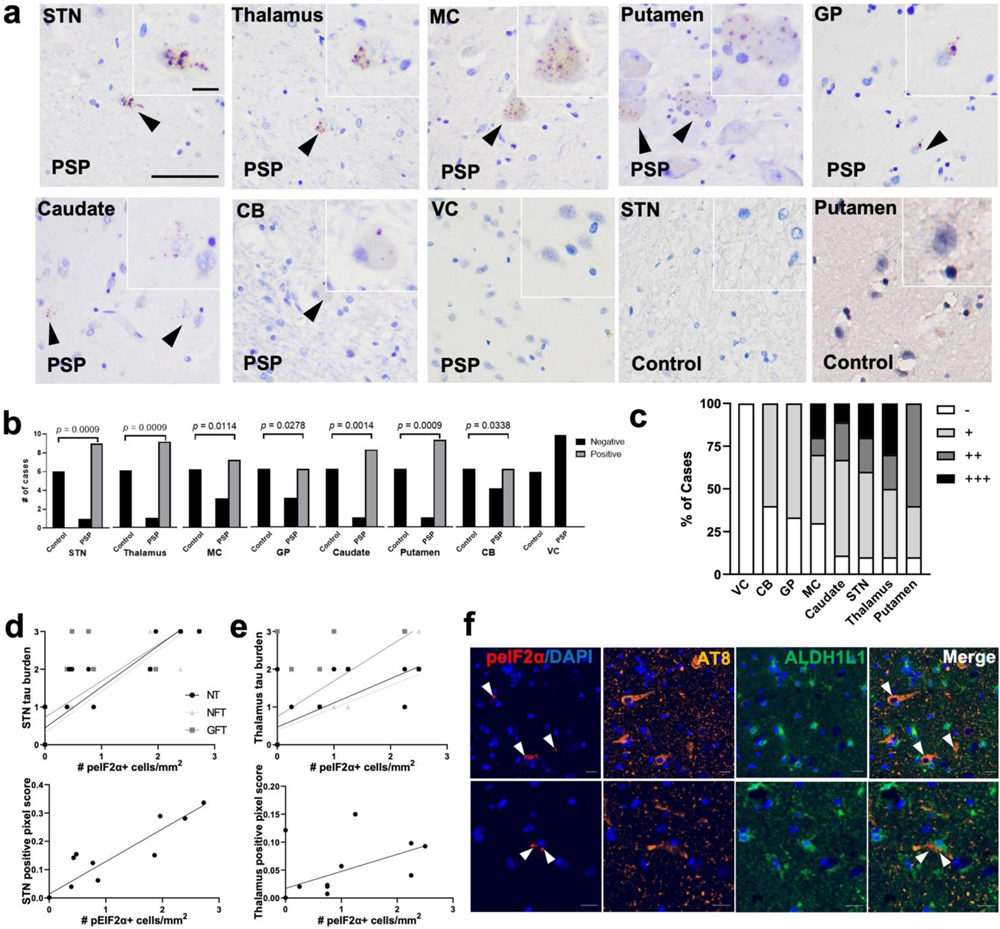
Elevated eIF2α and correlation with tau pathology in PSP vulnerable brain regions. **(a)** Representative images of peIF2α immunohistochemistry across brain regions showing peIF2α+ granules in all PSP regions except the visual cortex and no positive granules in controls, scale bar 50 µm to 10 µm. STN, subthalamic nucleus; MC, motor cortex; GP, Globus Pallidus; CB, cerebellum; VC, visual cortex **(b)** Quantification of (a) showing significant differences in the number of PSP and controls positive and negative for peIF2α+ cells in all brain regions except the visual cortex. Comparisons were performed using Fisher’s exact test. **(c)** Summary of scaled peIF2α burden across brain regions in PSP cases. Scaling was calculated using # peIF2α+ cells/mm^2^; -, 0; +, < 1; ++, < 2; +++, > 2. **(d-e)** Correlations between peIF2α and p-tau pathology in the STN (d) and thalamus (e), NT = neuropil threads; NFT = neurofibrillary tangles, GFT = glial fibrillary tangles. **(f)** Multiplex immunofluorescence showing colocalization of peIF2α (red) with p-tau (AT8, orange) positive neurons (top) and ALDH1L1 (green)-labeled astrocytes (bottom) in PSP, scale bar 25 µm.

**Table 4.** IHC Validation Patient Table.

|  | Group |  |  |
| --- | --- | --- | --- |
|  | PSP | Control | Total |
| Sample size, n | 10 | 6 | 16 |
| Avg age at death (range) | 70.8 (62.0-83.0) | 70.5 (62.0-82.0) | 70.7 (62.0-83.0) |
| Sex, male (%) | 70.00% | 83.30% | 75.00% |
| Average PMI (range) | 21.7 (10.6-37.8) | 22.9 (10.8 - 43.8) | 22.1 (10.6 - 43.8) |
| Brain weight (g) (range) | 1308 (1090-1577) | 1292 (1108 - 1380) | 1302 (1090-1577) |

To further investigate the relationship between increased EIF2 activation and tau accumulation, we explored whether tau burden correlated with the number of peIF2α^+^ cells in the subthalamic nucleus and thalamus. Phosphorylated tau burden was measured on whole slide images of AT8-stained sections by semi-quantitative scoring (-absent, + mild, ++ moderate, +++ severe) for neuropil threads (NT), neurofibrillary tangles (NFT), and glial fibrillary tangles (GFT), and by positive pixels. The number of peIF2α^+^ cells significantly positively correlated with all measures of tau burden in the subthalamic nucleus and thalamus (**Figure d-e, Table 5**).

**Table 5.**
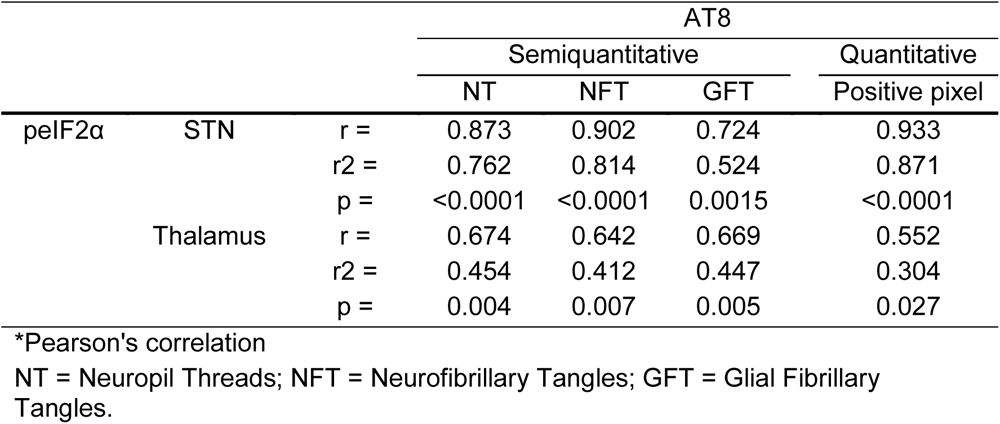
Summary of peIF2a and p-tau (AT8) correlation.

Finally, we performed multiplexed immunofluorescence to localize peIF2α positivity to different cell types by labeling nuclei (DAPI), astrocytes (ALDH1L1), peIF2α, and phospho-tau inclusions (AT8). We observed peIF2α granules in both p-tau positive neurons and p-tau positive astrocytes (**Figure 5f**). These results show EIF2 activation in PSP neurons and astrocytes with tau inclusions and provide histological validation to support the increased transcriptomic EIF2 activation observed in multiple cell types that are vulnerable to tau accumulation in PSP. Together these data show a quantitative association of peIF2α with tau burden in PSP specifically in vulnerable cell populations and brain regions.

### Differential gene expression and dysregulated functions in cell types not vulnerable in PSP

We also observed gene expression differences in other cell types not affected by tau accumulation in PSP (**Supplemental Figures 3**-**4**). PSP microglia overexpressed several ribosomal and heat shock genes related to topologically incorrect protein degradation, ER stress and PERK regulation as well as apoptosis genes such as *FOS* [74]. Microglia DEGs were associated with activation of EIF2 signaling and multiple cell death disease states (**Supplemental Figure 4b-e**). PSP endothelial cells differentially expressed several genes specific to endothelial cell function, homeostasis and secreted proteins including connective tissue growth factor (*CTGF)*, platelet endothelial cell adhesion molecule-1 (*PECAM1)* mitochondrially encoded cytochrome c oxidase (*MT-CO2)*, and metallothionein (*MT1X)* [75–77] (**Supplemental Figure 4b-c**). PSP ChP cells downregulated claudin-5 (*CLDN5*), the most enriched tight junction protein at the blood brain barrier [75] (**Supplemental Figure 4b-c**). Additionally, many PSP GWAS risk genes were overexpressed in cells not affected by abnormal tau accumulation (**Supplemental Figure 5**). *MAPT* was most highly expressed in microglia. *EIF2AK3, RUNX2*, and *DUSP1*, which is involved in the cellular response to environmental stress, were also highly expressed among microglia, OPCs, endothelial and ChP cells. This data suggests that PSP-related risk genes and cellular functions, such as *MAPT* expression, EIF2 signaling and ER stress, and blood brain barrier breakdown, are dysregulated in cell types not affected by p-tau accumulation and warrants further investigation into these cell types as reactive or promoting disease states in degenerating cell types in PSP.

## Discussion

These data demonstrate transcriptomic changes in all major cell types in the subthalamic region, which is selectively vulnerable in PSP, including cell types known to be susceptible to abnormal tau accumulation (i.e., neurons, astrocytes, and oligodendrocytes). Many genes related to protein translation and endoplasmic reticulum (ER) stress were differentially expressed, and EIF2 signaling was the top dysregulated pathway across vulnerable cell types. We performed histopathological validation, demonstrating co-localization of activated phospho-eIF2α (peIF2α) in p-tau positive neurons and astrocytes as well as a positive correlation of eIF2α activation with tau pathology burden. Brain regions that are most vulnerable in PSP displayed the highest peIF2α burden, whereas no signal was observed in protected regions. Additionally, we show distinct neuronal populations with differential gene expression patterns in PSP, some of which are reduced in PSP and enriched for apoptosis, and some that are not reduced in PSP and may reflect an adaptive response against neurodegeneration.

EIF2 signaling is activated by phosphorylation under stress conditions in a signaling program known as the integrated stress response (ISR) [21]. The ISR is activated by multiple cellular stressors, including heme depletion, ER stress, viral infection, and amino acid starvation [78]. ER stress is mediated by the unfolded protein response (UPR), which is composed of three branches initiated by protein kinase RNA-like ER kinase (PERK), inositol-requiring protein 1α (IRE1α), and activating transcription factor 6 (ATF6) [79]. GWAS studies identified *EIF2AK3*, which codes for PERK, as a risk allele for PSP, thereby implicating the PERK/EIF2 pathway, the ISR, and the UPR in the pathogenesis of PSP [17–19].

In this study, increased EIF2 signaling in PSP neurons was driven by altered expression of ribosomal, chaperone, and heat shock genes, as well as other genes related to protein quality control, the unfolded protein response and ER stress. Increased EIF2 signaling in PSP astrocytes and oligodendrocytes was driven by these gene sets and additionally by differential expression of genes specific to the individual UPR branches including PERK-regulated genes. UPR overactivation under chronic stress or abnormal eIF2α phosphorylation can trigger apoptosis [79, 80]. In our dataset, genes and pathways related to apoptosis regulation and autophagy signaling were also dysregulated in astrocytes, oligodendrocytes, and in the neuronal clusters with reduced proportions in PSP. Although the relationship between the UPR and autophagy has been well studied in neurons, its role in regulating cell autonomous and non-cell autonomous functions in other cell types has not been well studied [81–83]. Previous single-cell RNA-seq (scRNAseq) studies in tau burdened neurodegenerative postmortem brain tissue have identified similar stress response pathways in vulnerable cell types. In an AD scRNAseq study, AD astrocyte and neuron subclusters were enriched for genes relating to responses to incorrectly folded proteins [29]. A single-soma RNAseq study found that the ISR transcription factor ATF4, which is translated by eIF2α phosphorylation, and other heat shock proteins were highly upregulated in NFT-bearing neurons [31]. In a PD snRNAseq study, PD microglia and astrocytes upregulated chaperone and heat shock genes coupled with UPR dysregulation [34]. It is unclear if UPR-dysregulation is a common stress mechanism that affects multiple cell types in the brain during toxic misfolded protein aggregate-mediated neurodegeneration, or if the GWAS risk-associated *EIF2AK3* underlies a unique pathogenic mechanism specific to PSP.

UPR activation has been observed histologically in multiple neurodegenerative diseases, however, its role in disease pathogenesis is unclear. Here we show that EIF2 activation (peIF2α) in neurons and astrocytes positively correlates with p-tau burden and distribution patterns in brain regions vulnerable in PSP. Activated peIF2α co-localized with neurofibrillary tangles, and tau-positive astrocytes. Our study is the first to our knowledge to demonstrate histological evidence of UPR activation in astrocytes in postmortem tauopathy brain tissue. Other histological studies have also shown neuronal UPR activation occurs in PSP, AD, C9orf72-FTD, and PD post-mortem brain tissue [22–26, 84]. Neuronal PERK activation by phosphorylation (pPERK) is seen in PSP brain regions highly affected by tau and has been shown to co-localize with neuronal tau histologically [22]. However, other studies have found no correlation between tau accumulation and UPR activation in post-mortem AD and PSP brain tissue [85]. These previous studies have focused on total p-tau levels and neuronal p-tau, therefore it remains unclear if p-tau accumulation in glial cells induces PERK activation in human brain tissue.

*In vivo* studies have thus far not shown a consistent relationship between UPR activity and tau aggregation. Several studies have demonstrated UPR activation triggers tau accumulation and contributes to cell vulnerability and neurodegeneration. Increased UPR activation and vulnerability to ER stressors is seen in *MAPT*-mutation carrying iPSC-derived neurons with abnormal tau accumulation [86, 87]. Tau accumulation was shown to activate the UPR in human AD brains, tau^P301L^ mice and iHEK-Tau cells [88]. Activation of PERK-peIF2α-mediated UPR signaling in astrocytes *in vivo* and of mice infected with prion protein has been shown to drive non-cell autonomous neurodegeneration [89]. ISR activation in astrocytes was also shown to be an early, cell non-autonomous response to neuronal tau aggregation in a human neuron-astrocyte co-culture model [90]. Additionally, the UPR was identified as a top enriched pathway in a 3D neuron-astrocyte tauopathy assembloid model [91]. However, other studies found UPR activation to be neuroprotective and reduce p-tau burden. PERK activation reduced tau phosphorylation and increased cell survival in a 4R-tau overexpression neuronal model [92]. PERK activation also prevented tau aggregation *in vivo* and reduced PERK signaling correlated with increased tau neuropathology in postmortem AD brains [62]. Still other groups have found no correlation between tau accumulation and UPR activation in tau^P301L^ mice or tau^P301L^ primary neurons [93]. Many of these studies utilized mouse models or immortal cell line tau aggregation models to investigate UPR activation, which may not accurately recapitulate the cellular and molecular complexity of neurodegeneration in the human brain. Further examination of the mechanistic relationship between UPR activation and tau accumulation are needed in patient-derived induced pluripotent stem cell (iPSC) models that critically retain diseased patient genetic backgrounds.

Excitatory neuronal loss is a shared pathological feature across neurodegenerative disorders, including PSP. We identified two excitatory neuronal clusters, Ex1 and Ex3 with reduced relative proportions in PSP which are marked by apoptotic genes including *CELF2*. *CELF2* overexpression promotes apoptosis via regulating the endoplasmic reticulum associated degradation (ERAD) pathway and has been linked to AD [52, 94]. Ex1 and Ex3 cells from PSP patients overexpressed genes related to neuronal injury and downregulated several genes related to apoptosis regulation such as *JUND*, *IFI6,* and *MTRNR2L12. JUND* is proposed to protect cells from senescence and apoptosis, and *IFI6* is an interferon-stimulated gene that has been shown to inhibit apoptosis in vascular endothelial cells [95, 96]. *MTRNR2L12* codes for an isoform of humanin, a secreted peptide with neuroprotective effects by modulation of oxidative stress and apoptosis [97]. Humanin is protective against neuronal cell death in familial AD and *MTRNR2L12* is a potential blood biomarker for early AD-like dementia in Down Syndrome [98, 99]. Differential gene expression and pathway analysis indicated several critical neuronal functions including cAMP and calcium-mediated signaling are also dysregulated in these clusters. Disruption of calcium homeostasis and activation of calmodulin-dependent CAMKII has been shown to impair synaptic morphology and trigger neuronal apoptosis in AD [100, 101]. CAMKIIA also associates with and phosphorylates tau at multiple sites to induce a conformational change to generate paired helical fragment conformation [102–106]. Ex3 also upregulated calcium ion binding and calmodulin-mediated signaling regulator *PCP4.* Interestingly, PCP4 levels are decreased in the brains of patients with AD and HD, but not in PD [107]. Decreased expression of apoptotic regulating genes in the PSP-reduced Ex1 and Ex3 clusters may indicate neuronal populations vulnerable to tau accumulation or undergoing neurodegeneration.

We also identified two excitatory neuronal clusters, Ex2 and Ex4, that were not reduced in PSP. Both these protected neuronal populations are marked by *PP3CA*, *ARPP19* and *ARPP21*, genes related to protein phosphorylation, particularly tau phosphorylation. *PPP3CA* (also known as calcineurin) is the only phosphatase directly modulated by calcium to control synaptic transmission and other neuronal processes and is one of the major phosphatases that directly regulates tau phosphorylation and dephosphorylates p-tau in the brain [48–50, 108]. *PPP3CA* expression and activity is reduced in the brains of AD patients, and reduced *PPP3CA* activity leads to tau hyperphosphorylation in other tauopathies including chronic traumatic encephalopathy (CTE) [50, 109]. *ARPP19* and *ARPP21* are substrates for calcium dependent cAMP-dependent protein kinase (PKA), and expression of *ARPP19* is reduced in AD brains [110]. Reduced activity of the genes marking resistant clusters Ex2 and Ex4 are implicated in neurodegeneration, therefore the overexpression of these genes may indicate spared or protected neuronal populations. Of note, these cell clusters are mainly comprised of cells from samples with the largest cell counts.

Our data provide further evidence to support additional genes and mechanisms previously implicated in PSP, AD, and other neurodegenerative diseases, including iron accumulation and oxygen homeostasis, blood break barrier breakdown, and several AD-associated genes. PSP neurons and astrocytes overexpressed genes involved in heavy metal and iron accumulation and oxygen homeostasis, and ferroptosis signaling was overactivated in PSP astrocytes and oligodendrocytes. Multiple studies have shown ferritin protein defects, abnormal iron accumulation, and oxygen homeostasis in the brain is associated with neurodegenerative diseases, including PSP, PD, and AD [59, 60, 111–114]. A recent study showed iron accumulation and dysregulation of iron and oxygen homeostasis in astrocytes and oligodendrocytes in PSP postmortem brain tissue [60]. A single-soma RNAseq study found that iron homeostasis-associated genes *FTL* and *FTH1* were highly upregulated in NFT-bearing neurons [31]. We observed a low abundance of endothelial cells in PSP, which is consistent with the blood brain barrier breakdown seen in neurodegenerative diseases [115]. Several genes related to endothelial cell function and secreted proteins, including *MT1X*, were also differentially expressed in PSP endothelial cells, further suggesting blood brain barrier disruption. *MT1X,* which was downregulated in PSP endothelial cells, is involved in metal homeostasis and protection against oxidative stress [77]. PSP ChP epithelial cells downregulated *CLDN5*, the gene coding for the blood brain barrier tight junction protein claudin-5, whose dysfunction has been implicated in neurodegenerative disorders such as AD [75].

PSP astrocytes and oligodendrocytes also overexpressed many AD-associated genes, including apolipoprotein E (*ApoE*), clusterin (*CLU*), and *QDPR.* In an AD single-cell RNAseq study, *ApoE*, the most common risk factor for late-onset AD, was downregulated in AD OPC, oligodendrocyte, and astrocyte subpopulations, and upregulated in AD microglial subpopulations [29, 116]. Elevated clusterin levels are reported in AD-vulnerable brain regions and an AD-protective variant of *CLU* is associated with higher expression [117, 118]. Clusterin has been shown to be associated with tau in AD and primary tauopathies and is preferentially expressed by an AD pathology-associated astrocyte subpopulation [28, 119, 120]. Additionally, *CLU* loss is associated with exacerbated tau pathology in a tauopathy mouse model [120]. *QDPR* overexpression was previously identified in an AD-pathology associated white matter oligodendrocyte subpopulation [28]. PSP oligodendrocytes also had decreased expression of cholesterol-synthesizing *DHCR24.* Reduced expression of *DHCR24 also* occurs in the temporal cortex of AD patients [67]. Our PSP snRNA-seq data provides further evidence to support the role of iron accumulation and oxygen homeostasis, blood break barrier breakdown, and several AD-associated genes in neurodegenerative diseases.

In addition, we identified a novel astrocyte-oligodendrocyte hybrid cell population with activation of neuroprotective pathways. The hybrid cells were the only cell cluster to show significant downregulation of EIF2 signaling. PSP hybrid cells overexpress several genes that produce secretory proteins with known neuroprotective roles, including osteopontin (*SPP1*), carboxypeptidase E (*CPE*) and metalothionein2A (*MT2A*). *SPP1* has been shown to promote repair processes after injuries in the CNS by modulating inflammatory responses [121–124]. *CPE* is an extracellular neurotrophic factor that protects neurons *in vitro* from apoptosis and promotes neuronal survival [125]. Increased expression of metallothionein MT genes has also been shown in astrocyte subpopulations suggested to be neuroprotective and astroprotective in HD [30]. Increased metallothionein expression is associated with several neuronal injuries including ischemia, infection, HD, and AD [120]. Mice deficient in MT1/2 show reduced neuronal survival after ischemic injury, impaired wound healing and astrogliosis after freeze-injury [126, 127]. PSP hybrid cells also overexpress *S100A6* (calcylin), which marks astrocytic progenitors in the subgranular zone [128]. S100A6 is overexpressed by reactive astrocytes surrounding amyloid-β plaques in AD and neurodegenerative lesions in amyotrophic lateral sclerosis (ALS) [129–131]. S100A6 can bind to Zn^2+^ to prevent Zn^2+^ induced toxicity [132, 133]. Zinc can also bind tau directly to contribute to tau toxicity independent of tau hyperphosphorylation [134, 135]. S100A6 can also enhance the phosphatase activity of PPP5C towards p-tau [136]. Together, this data shows the novel astrocyte-oligodendrocyte hybrid cells overexpress neurotrophic factors in conjunction with suppression of EIF2-mediate stress response, suggesting a potential neuroprotective function or homeostatic role. The UPR maintains cellular homeostasis under conditions of protein aggregation and misfolding, therefore this cell state might also represent a cell intrinsic protective mechanism in the hybrid cells to prevent tau aggregation. Single-cell RNAseq studies investigating other neurodegenerative disorders such as AD and Huntington’s disease (HD) similarly identified hybrid cells which may represent intermediate cellular states [29, 30, 132]. However, the functional roles of these intermediate cellular states have not been elucidated. Injury and stress in the brain potentially drive molecular changes in certain cell populations to mitigate neuronal and glial damage. Future investigation will assess the potential functional role of the hybrid population.

Previous genetic studies have identified PSP susceptibility loci and candidate risk alleles, however one challenge in GWAS interpretation is identifying how genetic risk manifests across cell types [17–19]. We sought to address this by examining candidate risk gene expression patterns across different cell clusters utilizing an approach previously used in AD and PD single-cell RNAseq studies [29, 33, 34]. We observed PSP risk genes with both overlapping and divergent expression patterns in neurons, astrocytes, and oligodendrocytes. Our results also show that PSP risk genes are highly enriched not only in vulnerable cell types but also cells where toxic tau does not accumulate, such as microglia. *MAPT* showed coordinate downregulation in all PSP excitatory neuronal clusters, and only slight upregulation in oligodendrocytes as well as microglia. This supports previous studies suggesting that the PSP *MAPT-* related risk does not manifest as differences in total tau mRNA expression [137]. Although the *MAPT* locus is the most strongly associated genetic risk factor, the mechanism is still unclear. The PERK-encoding gene *EIF2AK3* was only upregulated in one neuronal cluster (Ex2). Tau aggregation and pharmacologic UPR activation *in vivo* altered the expression of PERK-regulated genes but not the expression of the PERK-encoding gene (*EIF2AK3)* [62]. This suggests that transcriptional changes during tau-induced UPR activation may be downstream of or not change *EIF2AK3* expression. *STX6* showed coordinated upregulation in all PSP excitatory neuronal clusters but not in glia, implicating a primarily neuronal *STX6*-linked risk. Genetic variation at *STX6* could influence movement of misfolded proteins from the ER to lysosomes via the endosomal system, and thus further implicate the UPR in PSP [17]. *MOBP*, a gene specific to oligodendrocytes, was downregulated in most cell types. Decreased *MOBP* expression could contribute to oligodendrocyte or myelin dysfunction in PSP [17, 19]. High expression of *SLC01A2* in PSP vulnerable cell types (neurons, oligodendrocytes, and astrocytes) suggests a potential coding variant link to tau accumulation. A variation in *SLCO1A2* has previously been associated with amyloid-β deposition in AD-related neurodegeneration [138]. Studies that examine the cell-type specific effects of PSP risk allele gene expression on neuronal, astrocyte, and oligodendrocyte function are needed. Together, these data reveal cell type-specific expression patterns of known PSP risk genes and warrant further examination in cellular systems, such as PSP patient-derived iPSC models of different cell types, which retain the diseased human genome.

A limitation of our study is the small sample size and heterogeneity of PSP pathology in patient samples. While this heterogeneity is likely representative of the spectrum of PSP disease severity, whether there are any significant associations of particular neuronal subpopulations with tau pathology requires systematic stratification of the PSP variants and further analysis with a larger cohort of individuals with similar PSP tau pathology. Further, the number of transcripts from PSP and control cases differed due to mitochondrial rate quality control processing. Without quality control processing, the number of transcripts from PSP cases was lower, however these may have arisen from the neurodegenerative changes that typify PSP, as PSP is a neurodegenerative disease with greater proportion of apoptotic cells than healthy control samples. To balance the number of detected genes, we applied an adaptive mitochondrial rate threshold that removes highly apoptotic cells while minimizing the loss on functional transcripts, however this may have had an effect on our results.

In conclusion, our study contributes a novel characterization of cell-type specific changes in PSP using snRNA-seq and provides transcriptomic and histological evidence for the involvement of adaptive stress pathways in PSP pathogenesis in vulnerable cell types. Understanding the differences in gene expression between affected and unaffected cell types may reveal molecular mechanisms underlying vulnerability and resistance to neurodegeneration. These results also highlight the need for detailed functional characterization of multiple cell types to develop better models and therapeutic modalities for PSP. Our data will inform future studies to identify the mechanisms of tau accumulation, degeneration, and repair in PSP.

## Acknowledgements

This work was supported by the Karen Strauss Cook Research Scholar Award to A.C.P and J.F.C, the National Institutes of Health (R01AG063819 and R01AG064020 to A.C.P, R01AG054008 and R01NS095252 to J.F.C, F32AG072837 to K.W. and NIA career development award P30AG066514), the Alzheimer’s Association (AARG22-973911 to A.C.P and AARF-22-974094 to K.W.), the Carolyn and Eugene Mercy Research Gift to A.C.P., the Alzheimer’s New Jersey to A.C.P, the Robert J. and Claire Pasarow Foundation to A.C.P., the Alexander Saint-Amand Scholar Award to J.F.C, and the Tau Consortium/Rainwater Foundation to J.F.C.

## Author contributions

A.S, A.C.P, K.W., and J.F.C conceptualized and designed the experiments. A.S generated the snRNA-seq data with support from K.F. W.S, A.S, K.W., M.M.K, and H.W. analyzed snRNA-seq data and visualized results. K.W, K.F, D.K.D., M.M.K., and T.C. generated neuropathological data and performed immunohistochemical analysis. B.Z provided data analysis tools. All authors read and approved the final version of the manuscript.

---

**Supplemental Figure 1.**
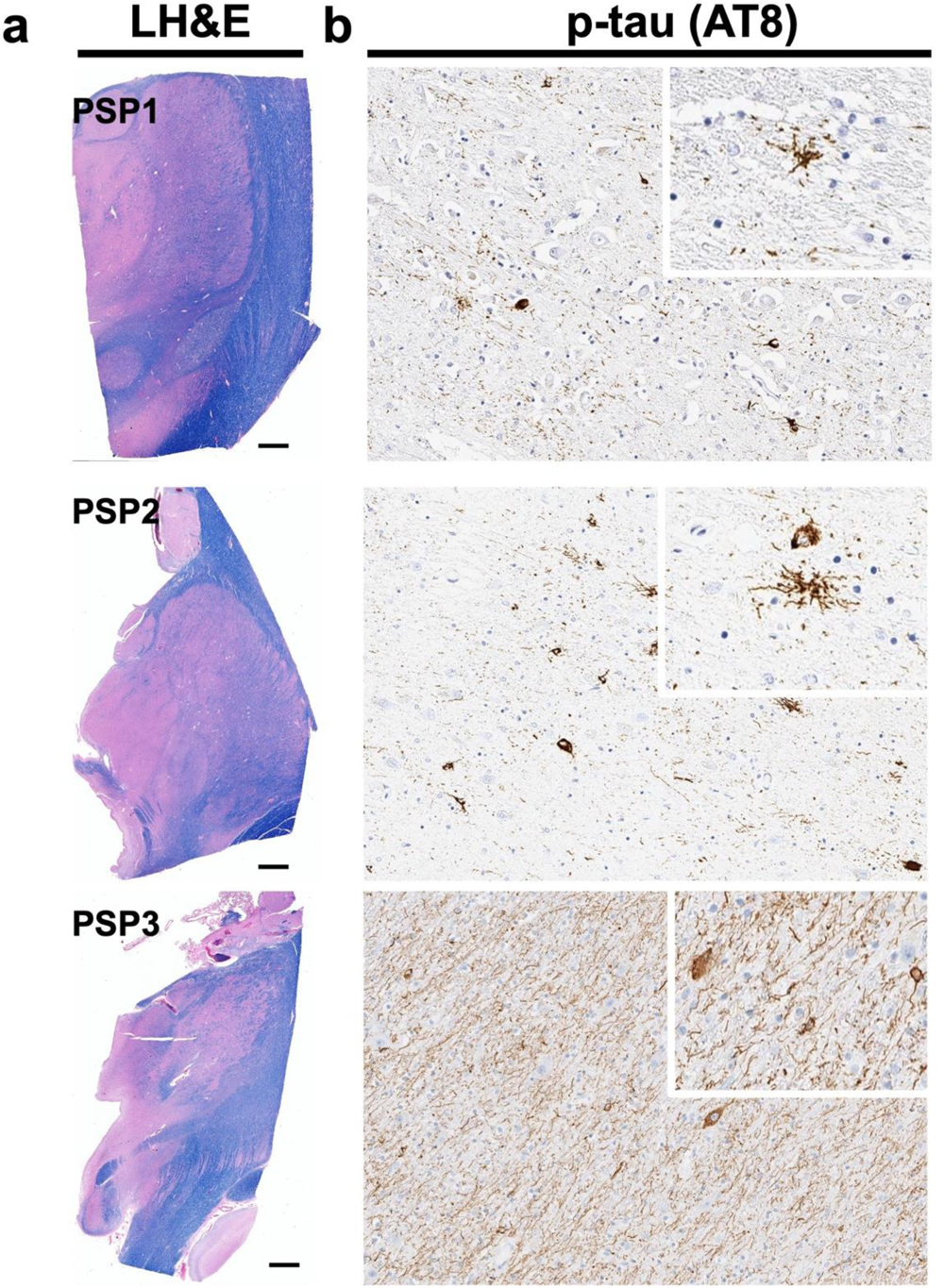
Tau burden and cellular pathology in PSP cases. **(a)** LH&E sections prepared from the contralateral hemisphere were screened for gross morphological characterization, scale bar 2 mm. **(b)** AT8 IHC shows varying p-tau neuronal, glial, and neuropil thread burden among the three PSP cases (20x).

**Supplemental Figure 2.**
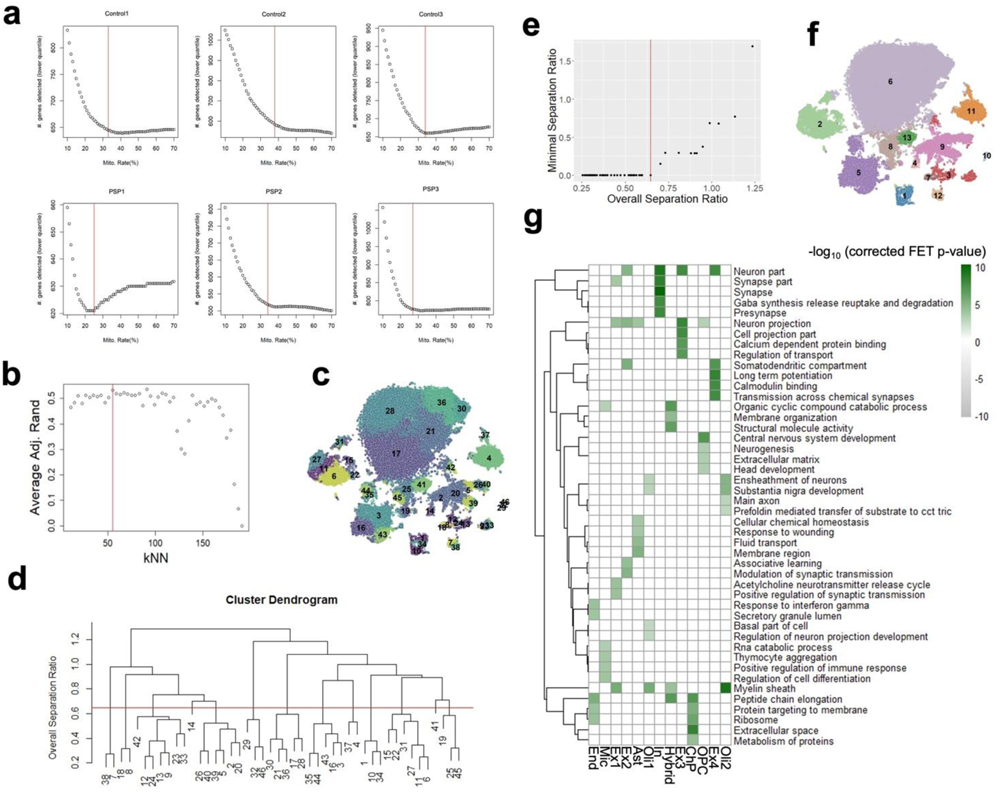
Adaptive mitochondrial thresholds per sample and identification of distinct cell clusters. **(a)**. Elbow plots showing the mitochondrial rate % per sample. Horizontal red lines mark the elbows as the final mitochondrial rate thresholds per sample. **(b)** Average adjusted Rand index of each clustering solution at different kNN parameters, compared to solutions at other kNN values. The red line remarks the optimal kNN value whose clustering solution shows the best overall adjusted Rand score. **(c)** tSNE plot of cells from MNN-corrected data. The clusters from the first-step of kNN clustering are labeled. **(d)** Dendrogram showing similarity between the initial cell clusters shown in (c), as evaluated by separation ratio. (**e)** Plot showing the overall separation ratio between all cell clusters merged in (d), calculated as the average separation between all clusters against the minimal separation ratio within clusters. The red line remarks the elbow point applied in (d). **(f)** tSNE plot of cells and cell clusters, refined by the separation ratio metrics in (d). Cluster identifications: 1 = End, 2= Mic, 3 = Ex1, 4 = Ex2, 5 = Ast, 6 = Oli1, 7 = In, 8 = Hybrid, 9 = Ex3, 10 = ChP, 11 = OPC, 12 = Ex4, 13 = Oli2. **(g)** GO enrichment heat map of the top five enriched functions and pathways per cluster in the cluster marker genes.

**Supplemental Figure 3.**
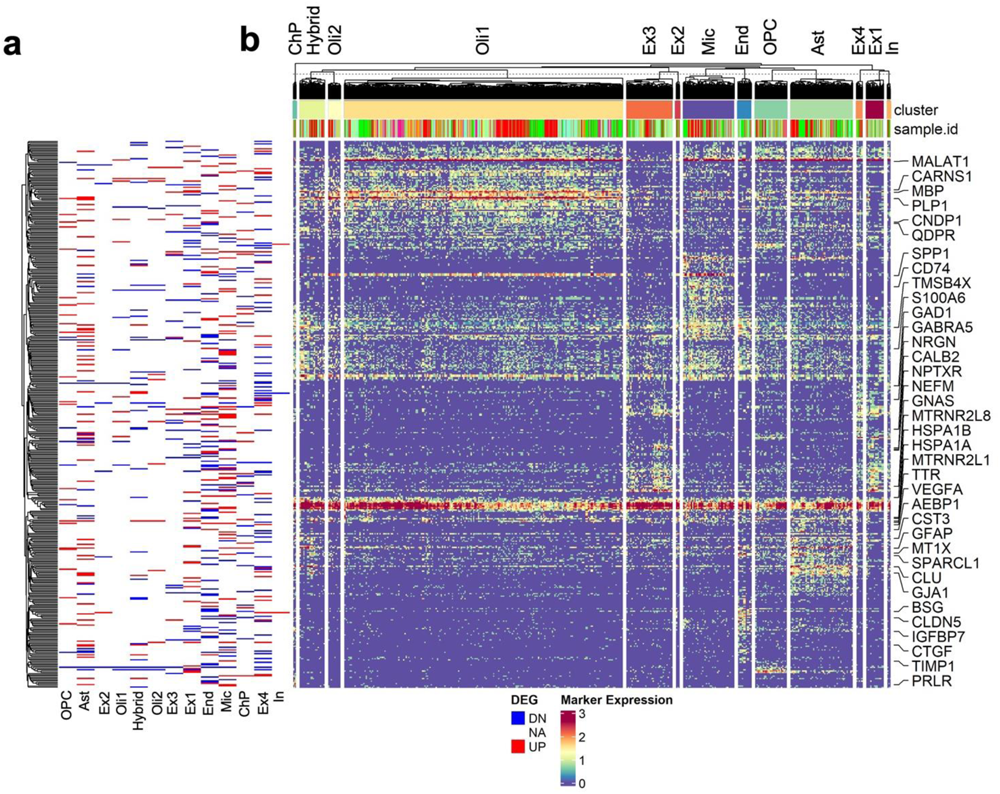
Differentially expressed genes (DEGs) across all cell clusters. **(a)** Tick graph showing the change direction of DEGs per cluster **(b)** Hierarchical clustering and gene expression heat map of log fold change of top DEGs (log fold change >1.2 and FDR < 0.05) between PSP and control cells across all 13 cell clusters.

**Supplemental Figure 4.**
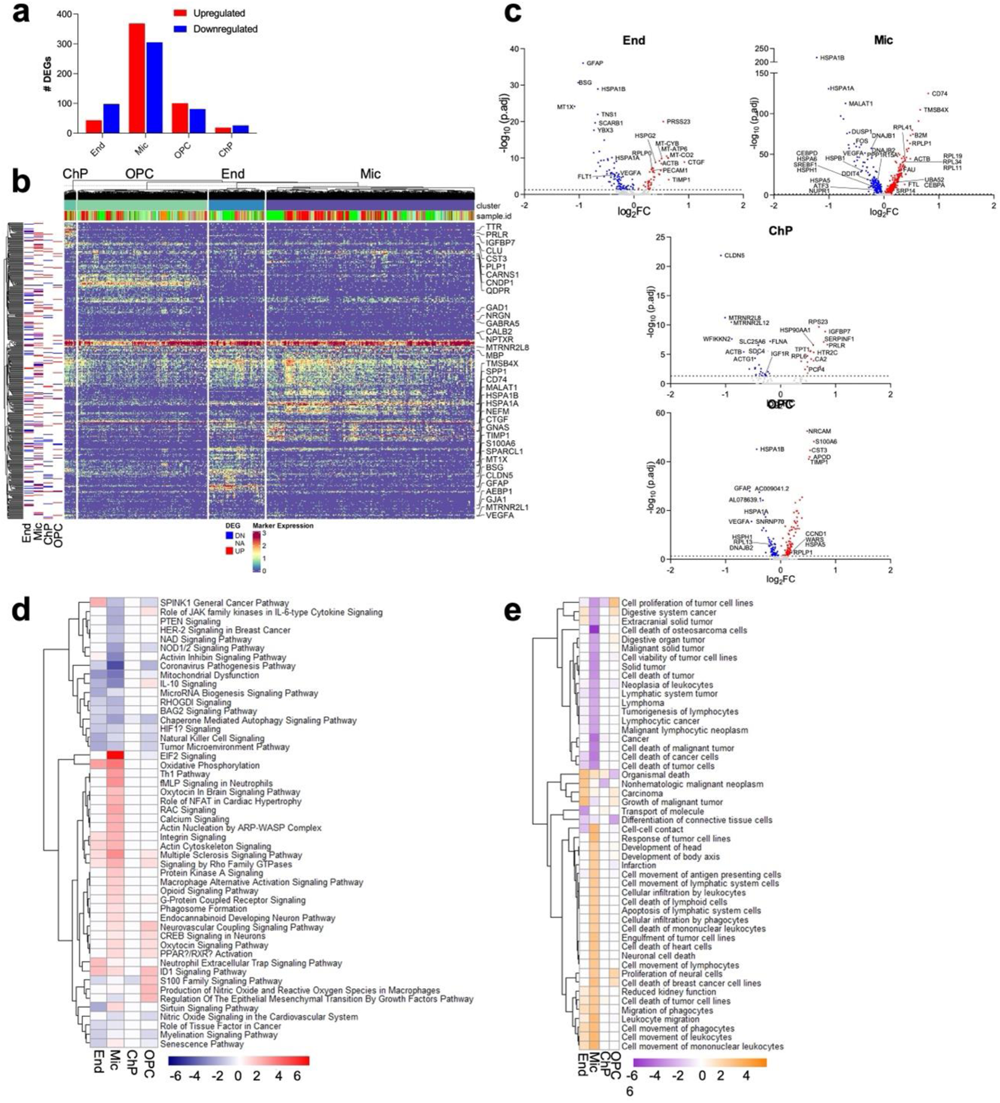
DEGs and regulated pathways in microglia, OPC, endothelial and choroid plexus clusters. **(a)** Total numbers of significant upregulated and downregulated genes in OPCs, microglia (Mic), endothelial (End) and choroid plexus (ChP) cells between PSP and controls. **(b)** Hierarchical clustering and gene expression heat map of log fold change of top DEGs (log fold change >1.2 and FDR < 0.05) between PSP and control cells. **(c)** Volcano plots showing select significant DEGs in individual cell clusters. **(d)** Top 50 differentially regulated pathways in PSP clusters derived from ingenuity pathway analysis (IPA). **(e)** Top 50 differentially predicted disease and function activation states in PSP clusters derived from IPA. For (d) and (e), p-values were generated from Fisher’s exact test and heat map is colored by z-score. Positive scores predict an overall increase in pathway activity or disease and function activation states and negative scores predict an overall decrease.

**Supplemental Figure 5.**
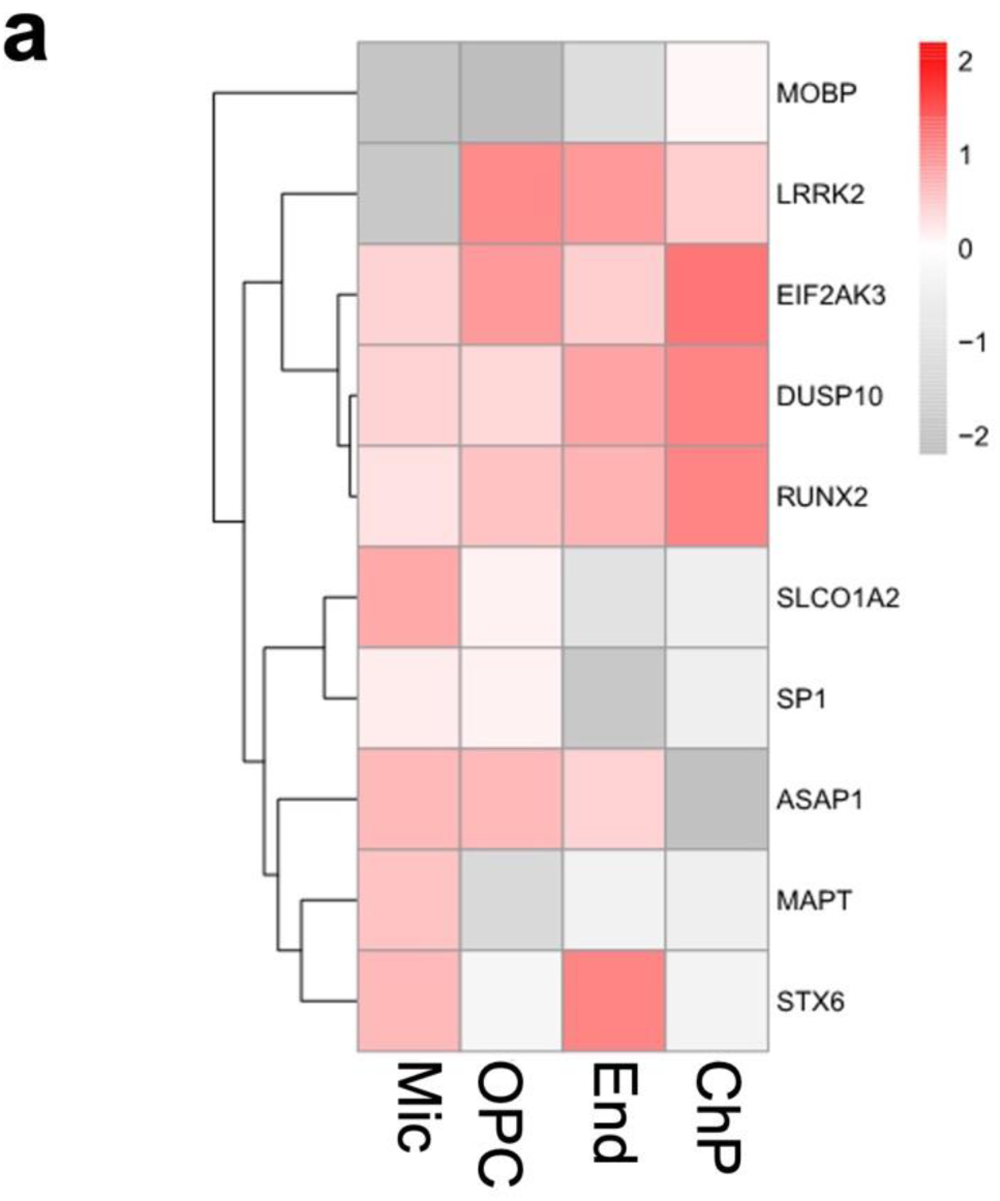
Expression patterns of PSP candidate risk genes across cell types not affected by tau accumulation in PSP. Heat maps showing the normalized log fold change of differential expression for candidate genes in PSP and control **(a)** microglia, OPC, endothelial and ChP cells.

